# Identification of novel human cellular substrates of *Staphylococcus aureus* serine protease SplB

**DOI:** 10.1101/2025.09.09.675177

**Authors:** Christopher Saade, Hannes Wolfgramm, Manuela Gesell-Salazar, Christian Hentschker, Katrin Schoknecht, Jens Fielitz, Britta Fielitz, Ning Li, Leif Steil, Uwe Völker, Barbara M. Bröker, Alexander Reder, Kristin Surmann

## Abstract

*Staphylococcus aureus* colonizes up to one third of the human population yet retains the capacity to cause invasive, life-threatening infections. The growing prevalence of antimicrobial resistance further complicates treatment. A major contributor to the versatility of *S. aureus* is its broad repertoire of virulence factors, among which secreted proteases facilitate dissemination from colonization sites into deeper tissues. Twelve extracellular proteases are secreted, with the serine protease-like protein (Spl) family (SplA–SplF), encoded within a single operon, accounting for half of them. Despite this prominence, the pathophysiological roles and substrate specificities of the Spl proteases remain poorly understood.

Here, we employed a direct protein–protein interaction approach to identify novel SplB substrates in human serum. We demonstrate that SplB cleaves three intermediate filament proteins, namely desmin, vimentin, and nestin, as well as heat shock protein β1 and α-enolase, which have not previously been recognized as targets of *S. aureus* proteases. Moreover, SplB was found to cleave native IgG, a feature otherwise described only for the glutamyl endopeptidase V8. These findings expand the host protein repertoire targeted by SplB and suggest broader roles for Spl proteases in immune evasion and tissue invasion.

## Introduction

*Staphylococcus aureus* (*S. aureus*) is a Gram-positive coccus belonging to the genus *Staphylococcus* (1). It colonizes approximately 25–30% of the human population (2), yet can rapidly switch into a virulent pathogen capable of invading and infecting virtually every tissue in the human body (3). *S. aureus* remains a leading cause of hospital-acquired infections and infection-related mortality (4), accounting for more than 1 million of the 3.7 million infection-related deaths worldwide in 2019 (5). Among the five leading pathogens, it is the only one responsible for over 1 million deaths annually (6).

In carriers, infections are typically caused by the colonizing strain (7), underscoring the ability of *S. aureus* to exist as a harmless commensal while retaining the potential to cause invasive disease. This remarkable versatility is largely attributed to its extensive network of virulence factors (8).

Extracellular proteases represent one class of virulence factors with crucial roles in bacterial dissemination, as well as immune modulation and evasion (9). *S. aureus* secretes twelve distinct extracellular proteases (10), of which six belong to the serine protease-like (Spl) family (11). Despite accounting for half of the secreted proteases, the physiological functions of the Spl proteases remain poorly understood.

All six Spl proteins (SplA–SplF) are encoded within a single operon (12). Their sequence identity ranges from 40% to 70%, with SplD and SplF sharing 95% sequence identity (13). Each Spl protease contains a conserved catalytic triad of serine, histidine, and aspartic acid (14). Functional studies suggest that Spls modulate the immune response by skewing T cells toward a type 2 phenotype, as indicated by elevated serum anti-Spl IgE and type 2 immune memory (15) (16). Other data suggest roles in bacterial dissemination, as Spls enhance bacterial spread in a rabbit pneumonia model (17). Nevertheless, their pathophysiological substrates remain largely unidentified.

Consensus cleavage motifs for Spl family members have been determined using peptide libraries and synthetic peptides, revealing distinct substrate preferences between Spls and suggesting specialized functions (18) (19) (20). Their narrow substrate spectrum supports a role in virulence and host–pathogen interactions rather than general protein degradation for nutrient acquisition. This is further reinforced by the coordinated regulation of the *spl* operon with other virulence determinants (13). Current evidence points to Spls as contributors to both dissemination and immune evasion.

Among the Spl family, SplB has been implicated in immune evasion through cleavage of complement system components, directly interfering with host defense mechanisms (21). SplB also targets ubiquitin-like modifying enzymes (22). Since the ubiquitin–proteasome system is increasingly recognized as an important factor in controlling intracellular *S. aureus*, SplB may influence host intracellular defense pathways (23).

Human serum represents a physiologically relevant environment containing a complex mixture of proteins under normal and pathological conditions (24). During bloodstream invasion, serum constitutes a hostile niche for *S. aureus*, with phagocytes, complement factors, and immunoglobulins contributing to antibacterial defense (25). This makes serum an ideal medium for identifying physiologically relevant protease substrates. However, the heterogeneity of serum proteins in size, chemistry, and post-translational modifications complicates labeling-based analyses (26). We, therefore, hypothesized that SplB targets serum proteins. To test this, we applied a systematic screen using a direct protein–protein interaction approach rather than labeling-based methods.

For this purpose, we employed the Strep–protein interaction experiment (SPINE), adapted from (27), which combines paraformaldehyde-mediated protein cross-linking with Strep-tag purification and mass spectrometry. Using this approach, we identified five previously unrecognized substrates of SplB in human serum. Furthermore, we demonstrate that SplB can cleave immunoglobulin G, a property previously described only for another *S. aureus* serine protease, the glutamyl endopeptidase V8 (9).

## Materials and methods

### Production of recombinant SplB wild-type and SplB mutant proteins

Recombinant production of SplB wild-type (SplB^WT^) and SplB mutant (SplB^Ser195Ala^), both with a C-terminal Twin-Strep-tag, was carried out as described elsewhere (28). In brief, *S. aureus* RN4220 was used to produce tagged SplB^WT^ and SplB^Ser195Ala^, which were encoded on the pTripleTREP plasmid (ProTec Diagnostics GmbH, Germany) (28). *S. aureus* was chosen as the expression host to preserve correct post-translational modifications and ensure proper signal peptide processing, both of which are critical for maintaining SplB enzymatic activity (19). Overexpression was induced with anhydrotetracycline (Merck KgaA, Germany), and the proteins were purified from the *S. aureus* culture supernatant via their C-terminal Twin-Strep-tag using affinity chromatography with StrepTactinXT according to the manufacturers protocol (IBA, Germany). The SplB^Ser195Ala^ variant was generated by substituting the catalytic serine with alanine in the SplB^WT^ sequence. Enzymatic activity was confirmed by incubating 2.5 µM SplB with 25 µM of the synthetic substrate 7-amino-4-methylcoumarin (AMC) coupled to Ac-VEID (PeptaNova, Germany), following the method described by Dubin and co-workers (19).

### SplB cross linking experiment in human serum

To identify SplB substrate candidates in human serum a Strep-protein interaction experiment (SPINE) was conducted as follows: 20 mL of human blood were collected in serum tubes (BD Biosciences, Germany), and serum was separated from whole blood by centrifugation at 1300 g for 8 min. The serum was diluted 1:20 in PBS (PAN Biotech, Germany) to a final protein concentration of 3.5 mg/mL and divided into three groups of samples, each containing four technical replicates (500 µL per replicate). No SplB was added to the first group, 10 µg/mL of SplB^WT^ were added to the second group, and 20 µg/mL of SplB^Ser195Ala^ were added to the third group. Samples were incubated for 5 min at 37 °C. For protein–protein cross-linking, a fresh solution from paraformaldehyde (Sigma-Aldrich, USA) was added to all groups simultaneously (final concentration of 0.25%) and incubated for 15 min at room temperature, followed by the addition of sarcosyl (final concentration of 1%) (Sigma-Aldrich, USA) in 50 mM HEPES buffer to adjust the pH of the protein mixtures to 8.0, optimizing conditions for subsequent affinity purification. Finally, 200 µL of MagStrep Type 3 XT Beads (IBA, Germany) were added to each sample to purify SplB via its C-terminal Twin-Strep-tag along with the covalently crosslinked serum proteins. The methylene crosslinks of the purified complexes were then hydrolyzed by an incubation at 95 °C for 2 h and then the proteins were digested using 160 ng trypsin/Lys-C (Promega, Germany) per 4 µg of protein and analyzed on an Orbitrap mass spectrometer (Thermo Fisher Scientific, USA).

### Identification of SplB substrate candidates

Substrate candidates were identified through a stepwise filtering and normalization strategy. To ensure data reliability, ions with a coefficient of variation (CV) greater than 0.9 across replicates were removed, thereby, eliminating highly variable signals. Only ions with a q-value ≤ 0.01 were retained to maintain a false discovery rate of 1% or lower. Furthermore, ions were required to be detected in at least 50% of biological replicates per condition, reducing the influence of sporadic identifications. Proteins were considered only if sequence coverage reached at least 10% and if they were supported by a minimum of two peptides, both of which are standard criteria for confident protein identification. For normalization, two reference proteins that showed medium intensity across all groups and the lowest standard deviation were selected as internal standards, and the intensity values of all other proteins were divided by these references. Based on these criteria, proteins were classified as potential SplB substrates if they were absent in the first group (no SplB), consistently detected in at least 50% of biological replicates in groups two and three, and enriched in group three relative to group two (ratio > 1.2).

### Protein cleavage assays

All substrate candidates were commercially available and purchased in recombinantly purified form: desmin (Progen, Germany), heat shock protein β1 (Creative Biomart, USA), α-enolase (Creative Biomart, USA), the constant domain of the immunoglobulin κ-chain (Antibodies-online.com, Germany), serpin A12 (Creative Biomart, USA), and human IgG isotype (Invitrogen, Germany) along with two additional proteins from the same family as desmin: vimentin (Thermo Fisher Scientific, USA) and nestin (Rockville, USA). All proteins were adjusted PBS to a final concentration of 1.5 µM and incubated under the following conditions: Either (i) alone for 24 h, or at a 1:4 enzyme-to-substrate ratio (ii) with SplB^WT^ for 4 h or 24 h, and (iii) with SplB^Ser195Ala^ for 24 h.

### Silver staining of SDS-PAGE gels

For the detection of cleavage bands, proteins were separated by SDS-polyacrylamide gel electrophoresis (PAGE) (NuPAGE™ 4 to 12 %, Bis-Tris, 1,0 mm, Midi-Protein-Gel, Thermo Fisher Scientific, USA). The gels were fixed in an aqueous solution containing 50% methanol (Merck KGaA, Germany), 12% acetic acid (Carl Roth, Germany), for at least 1 h or overnight. Gels were then washed two times with 50% ethanol for 10 minutes, briefly incubated for 1 minute in 0.02% sodium thiosulfate pentahydrate (Carl Roth, Germany). After two washing steps with distilled water, gels were stained with 0.2% silver nitrate (Sigma-Aldrich, USA). Lastly, gels were developed in a sodium carbonate solution (Merck KGaA, Germany) containing freshly added 37% formaldehyde (Sigma-Aldrich, USA) until protein bands became visible (4–10 min). Development was stopped by a 20-second incubation in 1% glycine (Carl Roth GmbH, Germany) followed by washing in distilled water for at least 15 min. Successful cleavage was confirmed by the presence of cleavage bands in protein samples incubated with SplB^WT^ and their absence in protein samples incubated alone or with SplB^Ser195Ala^.

### Western blot analysis

Proteins separated by SDS-PAGE were transferred onto PVDF membranes using the Trans-Blot Turbo system (Bio-Rad, Hercules, USA). Gels were equilibrated in transfer buffer (Bio-Rad, Hercules, USA), and membranes were briefly activated in 100% methanol (Merck KGaA, Germany) prior to assembly. The transfer was run at 1.3 A and 25 V for 7 minutes, followed by drying before downstream analysis. Prestained markers (Precision Plus Protein™ WesternC™, Bio-Rad, Hercules, USA), or PageRuler™ Prestained Protein Ladder (Thermo Fisher Scientific, USA) were used to confirm transfer efficiency. Membranes were activated in 100% methanol, blocked with Intercept® TBS (LI-COR Biosciences, USA), and incubated with primary antibodies α-SplB1 (29) (1:3000, mouse anti SplB monoclonal antibody produced in-house) or rabbit anti-desmin (1:60,000, Proteintech, USA). After washing with tris-buffered saline with tween 20 (TBS-T), they were exposed to secondary antibodies coupled with an infrared dye, including goat anti-mouse IRDye (1:1000), goat anti-rabbit IRDye (1:20,000) (LI-COR Biosciences, Lincoln, NE, USA), or labeled binder XT CW800 anti-strep tag (1:20,000). Post-incubation washes were followed by brief drying on Whatman® paper. Membranes were then exposed to the Odyssey® CLx imaging system (LI-COR Biosciences) with fluorescence detection in the 700 and 800 nm channels.

### High-efficiency Undecanal-based N Termini EnRichment (HUNTER) for cleavage site identification

Cleavage sites were identified using the HUNTER technique (30). Samples were divided into three groups: (1) substrate pool only, (2) substrate pool with SplB WT, and (3) substrate pool with SplB mutant (50 nM each in PBS). All measurements were conducted in three replicates. Substrates and enzymes were incubated at 37 °C for 24 h. The following day, 17 µL of each sample were reduced by addition of dithiothreitol (DTT, final concentration 10 mM; Sigma-Aldrich, USA) and incubation at 37 °C for 30 min. Resulting sulfhydryl groups were then alkylated by addition of iodoacetamide (IAA, final concentration 50 mM;Sigma-Aldrich, USA) and incubated at room temperature for 30 min in the dark. Excess IAA was quenched by addition of DTT (final concentration 50 mM; 20 min, RT, dark).

Isotopic dimethyl labeling was performed using light (^12^CH_2_, group 1), medium (^12^CD_2_, group 2), and heavy (^13^CD_2_, group 3) formaldehyde, each at a final concentration of 40 mM, in combination with light sodium cyanoborohydride (NaBH_3_CN, groups 1 and 2), and heavy sodium cyanoborodeuteride (NaBD_3_CN, group 3) at a final concentration of 20 mM (all Sigma Aldrich, USA). Samples were incubated at 37 °C and 700 rpm for 1 h. An additional labelling pulse was applied, followed by a second incubation as before. The labelling reaction was quenched with 2 M Tris (pH 6.8): Sigma-Aldrich, USA), to a final concentration of 600 mM and incubated at 37 °C for 3 h.

Protein cleanup and tryptic digestion were performed using SP3 beads (31) (hydrophilic and hydrophobic Sera-Mag SpeedBeads™ carboxyl magnetic beads mixed in a 1:1 ratio; Cytiva, Germany) in a protein-to-beads ratio of 1:5 (5 µg beads per 1 µg protein). Binding to the beads was performed in 80% ethanol shaking at 1500 rpm for 18 min. Bead-bound proteins were washed twice with 90% ethanol on a magnetic rack, then mixed with trypsin at a 1:25 enzyme-to-protein ratio. Samples were adjusted to 30 µL with 20 mM HEPES buffer (pH 8.0) and digested overnight at 37 °C.

After digestion, generated peptides were separated from the beads by centrifugation at 14,000 g for 1 min and incubation on a magnetic rack, followed by acidification to 1% acetic acid and pooling of samples from all three groups per replicate. Newly generated tryptic N-termini were hydrophobically labeled with undecanal, using undecanal in a 30:1 protein-to-undecanal ratio and NaBH_3_CN at a final concentration of 30 mM in 40% ethanol. The reaction was carried out at 37 °C with shaking at 800 rpm for 1 h. The undecanal-labeled tryptic peptides were depleted using ZipTip C18 tips (Sigma-Aldrich, USA). The flow through was subsequently lyophilized and reconstituted in 12 µL buffer A containing internal retention time (IRT) standard (1:50; Biognosys, Germany) for LC-MS analysis.

### MS measurement (HUNTER)

LC-ESI–MS/MS analyses of the obtained peptide solutions were conducted using a reverse phase HPLC chromatography system (Ultimate™ 3000 nano-LC system, Thermo Fisher Scientific) coupled to an Exploris™ 480 mass spectrometer (Thermo Fisher Scientific), in accordance with the settings outlined in the supplementary tables 1 and 2.

Database searches were MaxQuant (32). Databases contained the proteins of interest and the proteins of the production strains (*E. coli* K12). Semi N-terminal ArgC cleavage rules, 2 missed cleavages, fixed modification (cysteine carbamidomethylation), variable modifications (methionine oxidation, N-terminal acetylation) and the dimethyl labelling, N-terminal and at lysine, were considered.

We used R version 4.4.1 (33) and the following R packages: devtools v. 2.4.5 (34), ggrepel v. 0.9.5 (35), ggsci v. 3.2.0 (36), htmlwidgets v. 1.6.4 (37), openxlsx v. 4.2.6.1 (38), plotly v. 4.10.4 (39), tidyverse v. 2.0.0 (40).

### Identification of targets using SplB cross linking

A quantity of 4 µg of each sample was digested using a Trypsin/Lys-C Mix at an enzyme-to-protein ratio of 1:25 using the SP3 protocol (41). As previously outlined by Reder *et al*. (42), LC-ESI MS/MS measurements were conducted in accordance with the established protocol. In summary, the peptide mixtures were separated via nano-LC (UltiMateTM 3000) and analyzed on an Exploris™ 480 mass spectrometer (both Thermo Fisher Scientific, Bremen, Germany) in DIA mode (see also supplemental tables S1 and S4).

The data analysis was conducted using Spectronaut® (Biognosys AG, Zurich, Switzerland) version 18, employing the reviewed human database (version 2/2024, 20433 entries) provided by UniProt(43). Subsequent analysis was conducted utilizing R version 4.4.1 (44) and the SpectroPipeR package version 0.20 (45).

### Skeletal muscle myoblast cell differentiation to form myotubes

Primary human skeletal muscle myoblasts (HSMM; Lonza, Walkersville, MD, USA) were thawed and cultured according to the manufacturer’s instructions. Cells were maintained in Skeletal Muscle Growth Medium-2 (SkGM™-2 BulletKit™; Lonza, Walkersville, MD, USA), consisting of Skeletal Muscle Basal Medium-2 (SkBM™-2) supplemented with human epidermal growth factor (hEGF), dexamethasone, L-glutamine, fetal bovine serum (FBS), and gentamicin/amphotericin-B (SingleQuots™ Kit; Lonza, USA). Cultures were incubated at 37°C in a humidified atmosphere of 5% CO_2_. Medium was replaced every other day.

When cells reached 50–60% confluence, differentiation was induced by replacing SkGM™-2 with fusion medium consisting of DMEM:F12 (1:1; Lonza, Walkersville, MD, USA) supplemented with 2% horse serum (Lonza, Walkersville, MD, USA). The medium was exchanged every other day, and differentiation continued for 3-5 days until multinucleated myotubes were observed. For prolonged culture (>5 days), fusion medium was replaced with SkGM™-2 medium and refreshed every other day for up to two weeks.

#### Cell lysis

For downstream analysis, differentiated myotubes were lysed in radioimmunoprecipitation assay (RIPA) buffer (Thermo Fisher Scientific, Waltham, MA, USA) supplemented with protease inhibitor cocktail (Roche, Basel, Switzerland), when required. Lysates were collected and stored at −80°C until further use.

## Results

### Identification of SplB candidate substrates in human serum

The enzymatic activity of the C-terminally tagged recombinant SplB^WT^ protein was confirmed, while the corresponding SplB^Ser195Ala^ mutant form of the protease showed no activity. The assay was performed using the synthetic substrate 7-amino-4-methylcoumarin (AMC) coupled to the peptide Ac-VEID, as described by Dubin and co-workers (19). As expected, SplB^WT^ efficiently cleaved the synthetic substrate confirming its functionality, whereas no cleavage was detected with the mutant form (Supplementary Figure 1).

Next, applying the filtering criteria described in the methods section, we selected proteins that were (i) absent in group one (no SplB), excluding nonspecific binding to MagStrep Type 3 XT Beads, (ii) present in groups two and three and (iii) enriched by more than 1.2-fold in group three relative to group two, indicating specific enrichment in the presence of SplB^Ser195Ala^. This strategy yielded five substrate candidates: desmin (UniProt P17661), heat shock protein β1 (UniProt P04792), α-enolase (UniProt P06733), the constant domain of the immunoglobulin κ-chain (UniProt P01834), and serpin A12 (UniProt Q8IW75).

### Cleavage of the candidate substrates by SplB

To validate cleavage by SplB, the identified substrate candidates were purchased and incubated at an equimolar ratio (1.5 µM) with either SplB^WT^ or SplB^Ser195Ala^ for 24 h. The sodium/potassium-transporting ATPase subunit α-2 served as an internal specificity control. Four of the five candidates were cleaved by SplB^WT^ but left intact by SplB^Ser195Ala^ (Figure 1B). Only serpin A12, a serine protease inhibitor present in human serum and enriched by the SplB SPINE assay, showed no signs of cleavage similar to the control protein (Figure 1A). To further investigate the kinetics of cleavage of the identified substrates the four confirmed targets desmin, the constant domain of the immunoglobulin κ-chain (IGKC), α-enolase (ENOA), and heat shock protein β1 (HSPB1) were incubated under different conditions: alone (condition 0), with SplB^WT^ at equimolar ratio for 4 h (condition 1) or 24 h (condition 2), with SplB^WT^ at a 1:4 SplB^WT^:substrate ratio for 24 h (condition 3), and with equimolar SplB^Ser195Ala^ for 24 h (condition 4). Results are shown in figure 1.

**Figure 1:**
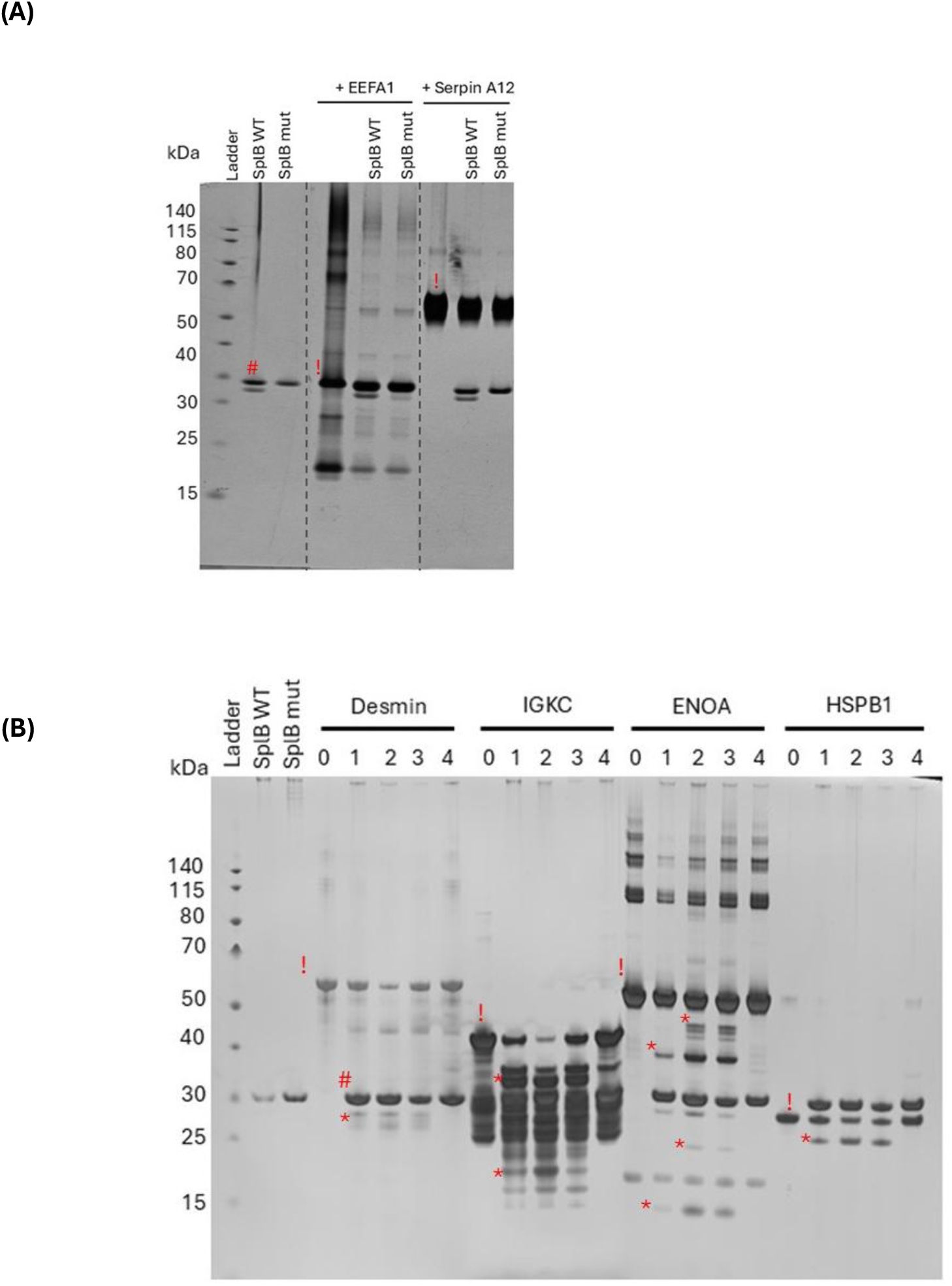
SplB_WT_ **cleaves desmin, the immunoglobulin κ-chain constant domain, α-enolase, and heat shock protein β1, but not serpin A12 or the sodium/potassium-transporting ATPase subunit α2**. **(A)** Silver-stained gel for serpin A12 and sodium/potassium-transporting ATPase subunit α-2 (EEFA1) incubated with SplB^WT^ or SplB^Ser195Ala^ (mut) at equimolar for 24 hours **(B)** Silver-stained gel of the *In vitro* cleavage products after incubating 1.5 µM substrate candidates with different concentrations of SplB^WT^ at various time points. Incubation with SplB^Ser195Ala^ served as control. Substrate candidates included desmin (∼55 kDa), recombinant IGKC (constant domain of the immunoglobulin κ light chain (∼12 kDa) fused to a glutathione S-transferase tag (GST; ∼26 kDa), α-enolase (ENOA∼ 49.3 kDa), and heat shock protein β1 (HSPB1∼ 28 kDa). The experiment was performed with n = 4 technical replicates, and a typical example is shown. Conditions included: 0: Substrate only (No SplB!); 1: SplB^WT^ and substrate at equimolar concentrations, 4-hour incubation; 2: SplB^WT^ and substrate at equimolar concentrations, 24-hour incubation; 3: SplB^WT^ 4:1 substrate-to-enzyme molar ratio, 24-hour incubation; 4: SplB^Ser195Ala^ and substrate at equimolar concentrations, 24-hour incubation. *= cleavage product != full length protein substrate #= SplB. Gel images were cropped for clarity, indicated by blue dotted lines; full gels are available upon request.

As shown in figure 1, the most efficient cleavage was observed for desmin and the immunoglobulin κ-chain constant domain, where cleavage products appeared as early as 4 h and almost no intact full-length protein remained after 24 h. Cleavage of α-enolase and heat shock protein β1 was slower: although cleavage products were also detectable already after 4 h incubation, a considerable fraction of full-length protein persisted even after 24 h with equimolar SplB^WT^. For heat shock protein β1, a single cleavage product was visible at 4 h, but only minimal progression was observed after 24 h.

### SplB cleaves IgG in its native form

The preparation of the recombinant fusion protein of domain of the immunoglobulin κ-chain with GST that was available for our cleavage assay with SplB, contained a considerable amount of contamination or degradation products that are visible in the control conditions (0 and 5). Since the aim of this study was to identify physiologically relevant substrates of SplB, we next tested whether SplB can cleave IgG in its native form. For this, a solution of human IgG was incubated with SplB under the same conditions described above. As shown in Figure 2, SplB cleaved IgG in its native form, producing one major and one faint band after 24 h of incubation at an equimolar ratio. A faint cleavage band was also detectable at a 1:4 SplB:IgG ratio, indicating that SplB can indeed cleave native IgG, although to a lesser extent than observed with the recombinant immunoglobulin κ-constant chain.

**Figure 2:**
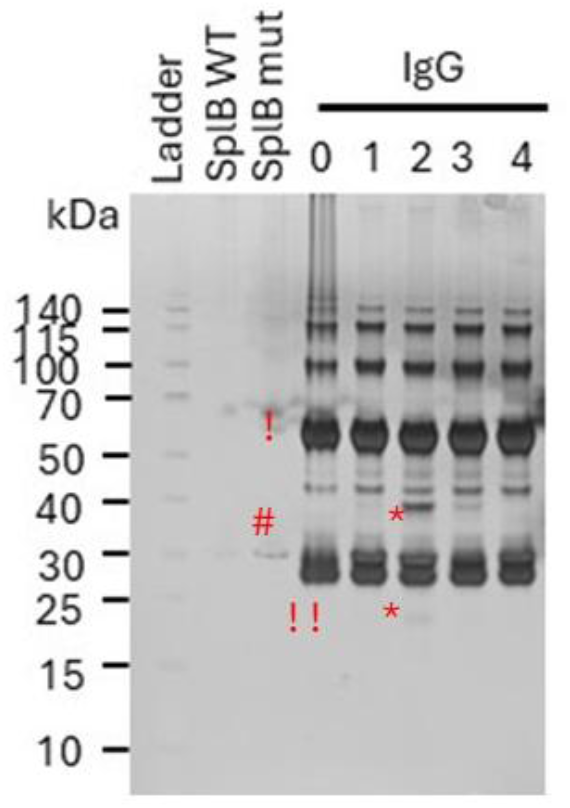
SplB cleaves immunoglobulin G. *In vitro* cleavage products after incubating 1.5 µM of whole IgG isotype solution with different concentrations of SplB-^WT^ at various time points, alongside SplB^Ser195Ala^. Reduction with β-mercaptoethanol resulted in the separation of IgG into its constituent chains. Two major bands are visible: the heavy chain at approximately 50 kDa and the light chain at approximately 25 kDa. The experiment was repeated with n = 2 technical replicates. Conditions included: 0: Substrate Only (No SplB); 1: SplB^WT^ and substrate at equimolar concentrations, 4-hour incubation; 2: SplB^WT^ and substrate at equimolar concentrations, 24-hour incubation; 3: SplB^WT^ 4:1 substrate-to-enzyme molar ratio, 24-hour incubation; 4: SplB^Ser195Ala^ and substrate at equimolar concentrations, 24-hour incubation. *= cleavage product, != IgG heavy chain, !!= IgG light chain, #= SplB

### SplB cleaves the intermediate filament proteins vimentin and nestin

The identification of desmin, a type III intermediate filament (IF) protein, raised the question of whether SplB can also cleave other members of the IF protein family. To address this, we tested two additional IF proteins: vimentin (type III) and nestin (type VI), both purchased in recombinant form and incubated with SplB under the same conditions as described above.

As shown in figure 3, vimentin was cleaved by SplB with the highest efficiency among all substrates tested. Complete degradation of full-length vimentin was observed after only 4 hours of incubation with SplB^WT^, and no intact protein remained under any of the tested conditions. In the case of nestin, the full-length protein could not be clearly visualized under the applied conditions due to its large size (>170 kDa). Nevertheless, multiple cleavage products appeared consistently in the presence of SplB^WT^, with several bands detected between 40–70 kDa and additional fragments around 25 kDa. These results suggest that SplB is capable of cleaving nestin at multiple sites, although the overall efficiency cannot be directly compared to the other identified substrates.

**Figure 3:**
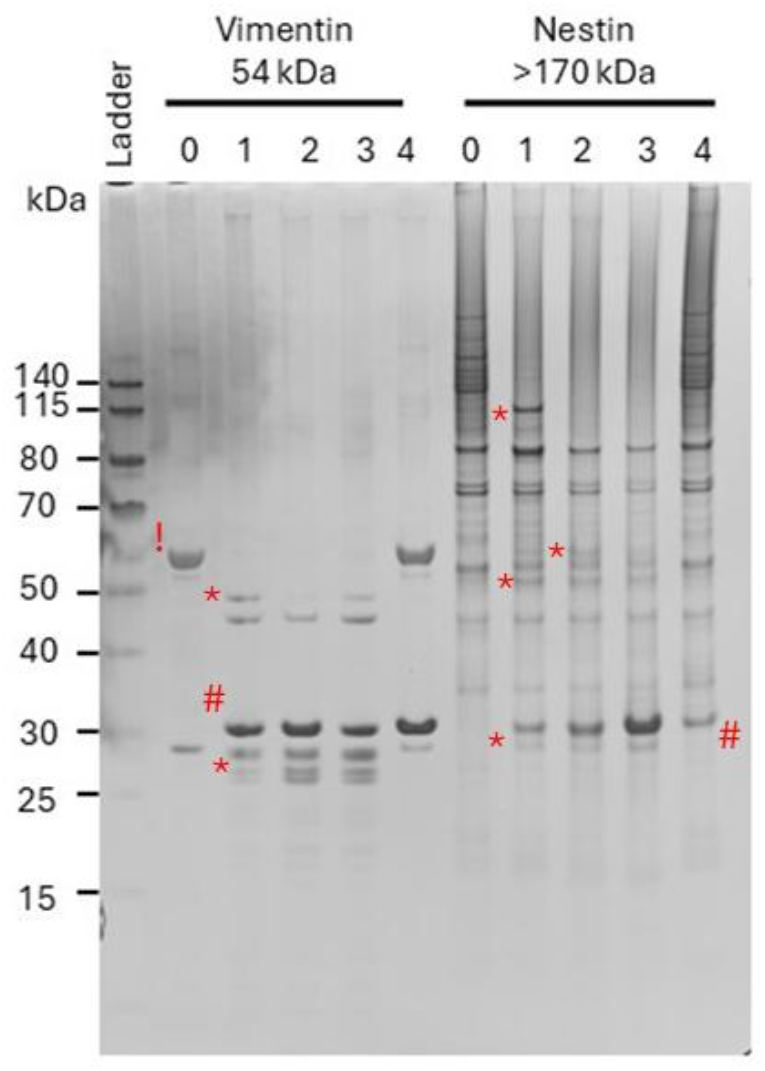
SplB cleaves the intermediate filament proteins vimentin and nestin. *In vitro* cleavage products after incubating 1.5 µM of vimentin (∼54 kDa) and nestin (>170 kDa) with different concentrations of SplB^WT^ at various time points. SplB^Ser195Ala^ served as control. The experiment was performed with n = 3 technical replicates, a typical gel is shown. Conditions included: 0: No SplB; 1: SplB^WT^ and substrate at equimolar concentrations, 4-hour incubation; 2: SplB^WT^ and substrate at equimolar concentrations, 24-hour incubation; 3: SplB^WT^ 4:1 substrate-to-enzyme molar ratio, 24-hour incubation; 4: SplB^Ser195Ala^ and substrate at equimolar concentrations, 24-hour incubation. *= cleavage product != full length protein substrate #= SplB.

### SplB cleaves native human desmin

Desmin is a muscle-specific type III intermediate filament that is essential for maintaining proper muscular structure and function (46). To assess SplB activity on endogenous desmin, we prepared cell lysates from skeletal muscle myoblasts differentiated into myotubes, as desmin is prominently expressed in skeletal muscle cells. First, the desmin abundance in the lysate was estimated by Western blotting, then equimolar amounts of SplB^WT^ or SplB^Ser195Ala^ were added and samples were incubated for 24 h (Figure 4). Western blot analysis showed that SplB^WT^ efficiently cleaved the native desmin in the myotube lysate: after 24 h, full-length desmin was no longer detectable, and a small cleavage fragment appeared just below the SplB band, mirroring the pattern observed with recombinant desmin in silver-stained gels. In contrast, no degradation was detected in any condition containing the catalytically inactive SplB^Ser195Ala^. Consistent with this, recombinant desmin was also fully cleaved after 24 h by Spl WT but not by Spl mut.

**Figure 4:**
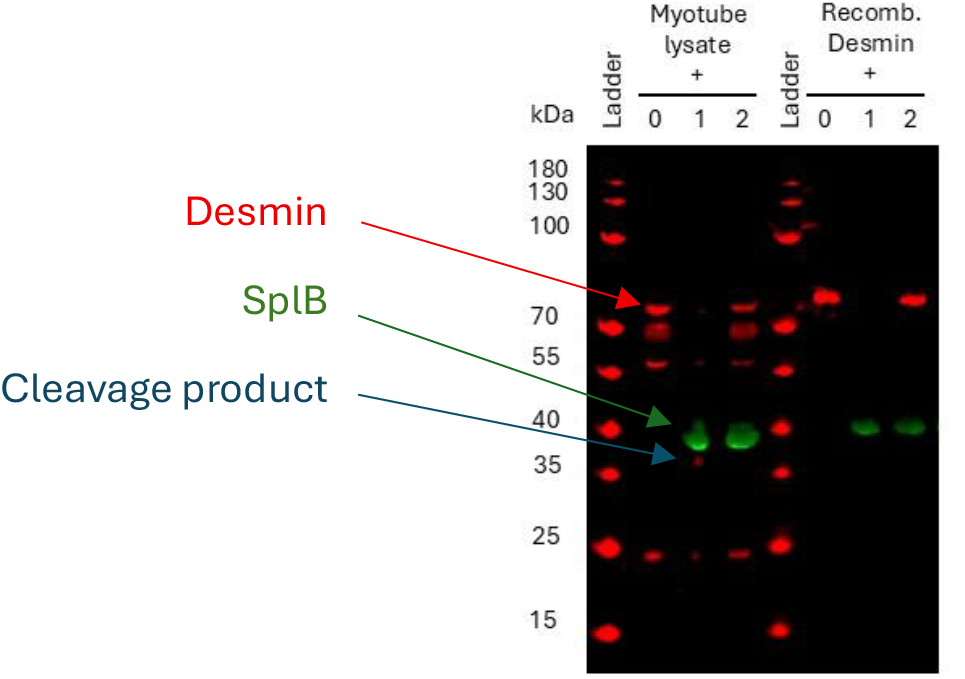
SplB cleaves endogenous human desmin. Cell lysates of human myotubes were prepared and the ability of 47.2 ng of SplB^WT^ to cleave 100 ng of native and recombinantly expressed desmin was assessed after 24 h of incubation. Conditions included: 0: Lysate or substrate only (No SplB); 1: SplB^WT^ and substrate at equimolar concentrations, 24-hour incubation; 2: SplB^Ser195Ala^ and substrate at equimolar concentrations, 24-hour incubation. A Western blot was performed using primary polyclonal anti-desmin antibodies (rabbit), detected with secondary goat anti-rabbit RD680 antibody (red color). A fluorescently labeled TactinXT CW800) (green color) was used to visualize the Twin-Strep tagged SplB proteins.

### SplB cleaves after glutamine (Q) and rarely after glutamate (E) at P1

To identify the SplB cleavage sites in the newly identified substrates, we applied the label-based HUNTER approach. Pools of recombinant substrates (50 ng each) were incubated for 24 h under three conditions: substrate alone, substrate with SplB^WT^ (50 ng), and substrate with SplB^Ser195Ala^ (50 ng). Each condition included three technical replicates. Stringent criteria were applied to confidently identify enzymatic cleavage sites: only N-terminal peptides that showed more than twofold enrichment in both comparisons, SplB^WT^ versus no SplB, and SplB^WT^ versus SplB^Ser195Ala^, were considered as cleavage products of SplB. Alignment of these peptides with the protein sequence of the substrate revealed the cleavage site. As shown in figure 5, no cleavage site could be determined for α-enolase with confidence, as none of the enriched peptides passed both criteria. In contrast, at least one highly confident cleavage site was identified in each of the other substrates. Desmin was cleaved at Q133, while heat shock protein β1 revealed a cleavage site after Q at position 176. Vimentin was cleaved after Q at position 82. For nestin, multiple cleavage sites were detected, with the strongest enrichment of new N-termini after Q at positions 33, 1239, 1491, and 1560, along with one site after E at position 1231, consistent with the multiple cleavage fragments observed in the silver-stained gels.

**Figure 5:**
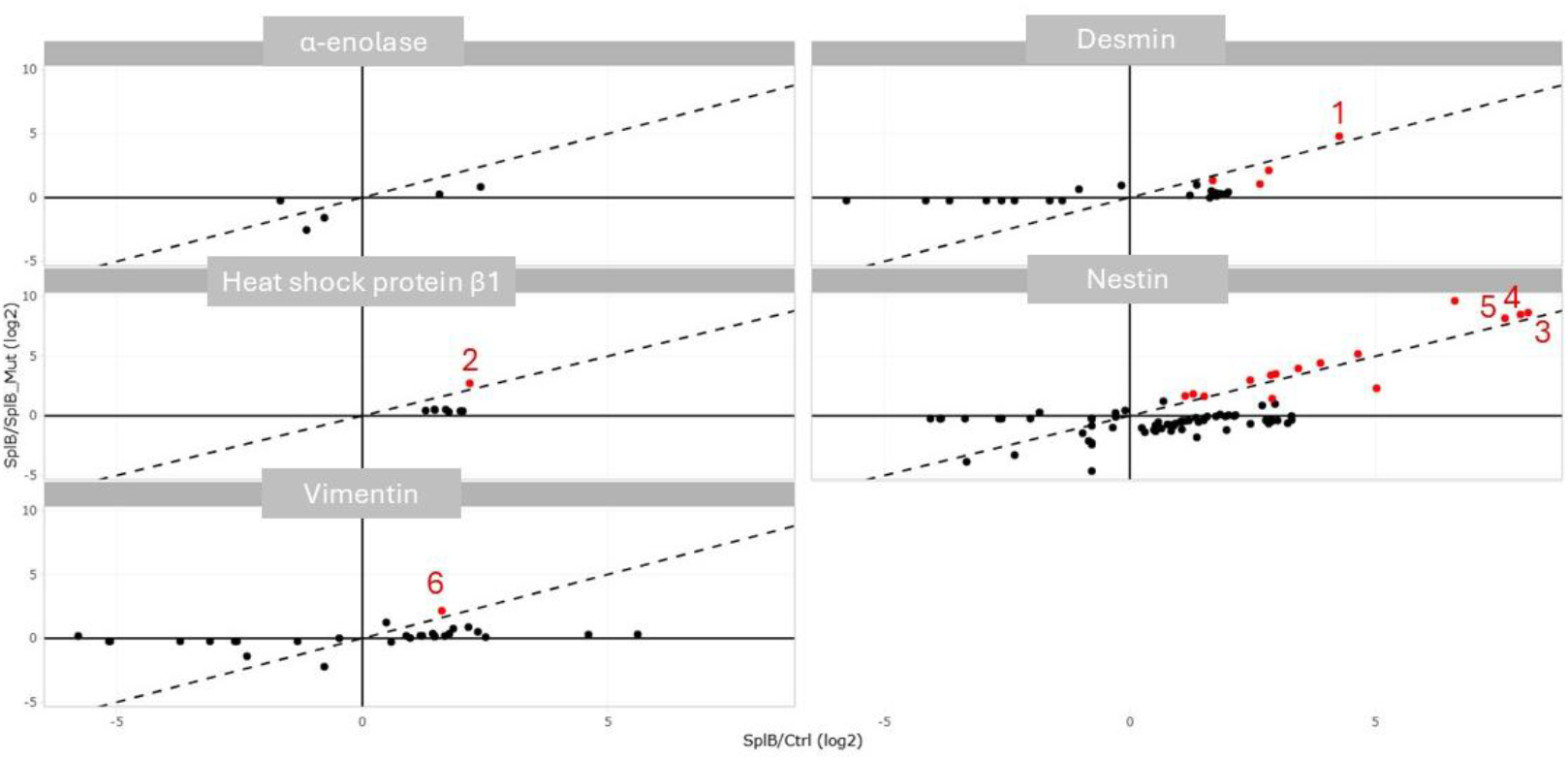
SplB cleaves after glutamine or glutamic acid. HUNTER analysis was used to identify SplB cleavage sites by comparing N-terminal peptide enrichment across conditions. The x-axis shows peptide enrichment in SplB^WT^–treated versus untreated substrate pools, and the y-axis shows enrichment in SplB^WT^ versus SplB^Ser195Ala^ (≥2-fold). Data represents three biological replicates. Red dots mark peptides ≥2-fold enriched in both comparisons, indicating specific SplB^WT^ cleavage sites. Identified sites included (1) desmin (Q133), (2) heat shock protein β1 (Q176), (3,4,5) nestin (Q33, Q1560, Q1491), and (6) vimentin (Q82). Corresponding peptide sequences were aligned to the protein sequence; the P1 residue (N-terminal of the peptide) was deduced and found to be Q or E. Sequences C-terminal of the cleavage sites, containing residues at P1′, P2′, P3′, etc., are shown.

## Discussion

In this study, we used a direct protein–protein interaction approach to identify candidate substrates of the secreted *S. aureus* serine protease SplB in the complex protein mixture of human serum. We adapted the SPINE method described by Herzberg and colleagues (2007) (27), using paraformaldehyde as a crosslinker and a C-terminal Twin-Strep tag for affinity purification of the SplB together with the covalently bound substrate proteins for identification by mass spectrometry. This approach has proven to be highly effective. Our criteria for candidate substrate detection were very stringent, allowing us to rapidly narrow down potential targets. Among the five enriched candidates, four were validated as true substrates. As expected serpin A12, a serine protease inhibitor (47), was enriched in the SplB complex but not cleaved in the follow up *in vitro* degradation assays. The identification of this inhibitor represents an ideal positive control, as its enrichment directly reflects the inhibitory interaction with SplB. This finding underscores the strength of our approach, which not only enables the detection of true substrates but also uncovers likely physiological inhibitors, thereby, providing insights into potential host defense mechanisms against pathogenic proteases. Moreover, the use of the catalytically inactive SplB^Ser195Ala^ instead of the WT protease proved to be particularly effective for substrate identification. While the wild-type enzyme can bind to substrates and subsequently cleave them, the inactive mutant remains bound without catalyzing proteolysis. This preserves the integrity of the interaction, thereby facilitating re-purification and detection of bound substrate candidates. Consistent with this, Consistently, SplB^WT^ enriched only serpin A12, whereas all validated substrates were enriched with SplB^Ser195Ala^ form.

One inherent limitation of this approach is the potential enrichment of biotinylated or otherwise surface-reactive proteins that bind nonspecifically to the StrepTactinXT beads used for purification of the tagged protein complexes. Nevertheless, such biotinylated proteins can be efficiently eliminated by pre-incubation with avidin, thereby, reducing background enrichment and increasing assay specificity.

Several components of the complement system such as C2, C6, and C8, which have previously been described as SplB substrates (21), were enriched in the presence of SplB with a fold change above 1.2. However, these proteins were also detected in the control group in the absence of SplB, suggesting nonspecific bead binding. This is likely explained by the intrinsic properties of complement proteins: they are abundant serum proteins with high surface affinity, prone to nonspecific adsorption due to hydrophobic or electrostatic interactions (48). Moreover, complement factors such as C3 and C4 contain reactive thioester groups that can form covalent linkages with exposed surfaces upon spontaneous or *ex vivo* activation, further contributing to their unintended enrichment (49). Thus, while true complement substrates of SplB may exist, careful discrimination from nonspecific background binding is essential in proteomic screens based on affinity purification.

The IF family protein desmin was identified as a SplB substrate, with both recombinant and native desmin cleaved in a similar manner. This was unexpected, as desmin is generally considered an intracellular protein (46). Its detection in serum likely reflects minor cell lysis *in vivo* or during blood handling. The specific enrichment of desmin among SplB-bound proteins highlights the sensitivity of our approach to detect potential substrates even at low abundance. IF proteins share a conserved central α-helical rod domain that mediates coiled-coil dimer formation, flanked by variable N-terminal head and C-terminal tail domains that differ among family members and determine tissue-specific functions (50). Extending this analysis, we found that SplB also cleaves two additional IF proteins, vimentin and nestin. Structural mapping revealed that desmin cleavage at Q133 lies at the junction of coil 1A (residues 109–141) and linker 1 (residues 142– 151) within the rod domain. Vimentin cleavage occurred at Q82 within its head domain, whereas nearly all identified nestin cleavage sites were located in its exceptionally long tail domain. These domains (head, linker, and tail) are long, disordered, and exposed, which may explain the higher cleavage efficiency observed for nestin compared to vimentin and desmin. Notably, cleavage of vimentin within its head domain is also mediated by the Chlamydia trachomatis proteasome-like activity factor (CPAF) to remodel host IFs and evade cytoplasmic immune surveillance (51). Previous work has further demonstrated that Spl promotes bacterial dissemination in rabbit lungs (17). Our finding that SplB efficiently cleaves IF proteins, including vimentin, suggests a potential mechanism for this observation, as lung tissue is particularly rich in IF proteins such as vimentin and cytokeratins (52). Consistent with the high degree of sequence homology among human, mouse, and rabbit IFs, our unpublished data show that SplB also cleaves mouse desmin in lung lysates.

The small heat shock protein β1 was another novel substrate of SplB. The SplB mediated cleavage happened at Q176 at C-terminal long disorded exposed tail. The heat shock protein β1 (also known as Hsp27 or HSPB1) is a stress-induced chaperone that regulates protein folding and IF stability (53)(54). The restricted cleavage pattern we observed may reflect functional interference with stress recovery rather than complete protein degradation. As HSPB1 dysfunction has been linked to desmin-related myopathies (55), its cleavage by SplB could synergize with IF disruption to compromise host cell integrity. Together, these findings suggest that SplB-mediated targeting of IF and the heat shock protein β1 may play an important role during the intracellular phase of *S. aureus* infection, where disruption of IF scaffolds could facilitate host cell damage and bacterial dissemination.

SplB also cleaved α-enolase, a glycolytic enzyme with critical immunological functions (56)(57). Because it supports dendritic cell activation (58), T cell proliferation (59), and plasminogen binding (60)(61), its cleavage could attenuate both innate and adaptive immune responses. Adding another substrate for SplB that is central for proper immune response.

Finally, SplB was able to cleave native IgG, although less efficiently than the recombinant κ-light chain–GST fusion protein. Native IgG produced fewer and weaker cleavage products, likely due to structural constraints of the antibody or altered accessibility introduced by the GST tag. SplB generated a dominant cleavage fragment with some evidence of light and heavy chain involvement, indicating limited activity against intact antibodies. This suggests that SplB may contribute to immune interference not only through intracellular regulators but also by targeting extracellular antibody structures, a mechanism reminiscent of other *S. aureus* proteases such as V8 (62).

Interestingly, several SplB substrates, including desmin, vimentin, and nestin, are primarily IF proteins, while the heat shock protein β1 and α-enolase also have major intracellular roles. Along with the identification of ubiquitin-modifying enzymes (22), this suggests that SplB may act during the intracellular phase of *S. aureus* infection. *S. aureus* can be internalized by professional phagocytes as well as non-phagocytic cells (63), such as epithelial and endothelial cells and osteoblasts (64), where it can escape the phagosome and replicate in the cytoplasm (65), coming into contact with IF. Cleavage of these cytoskeletal and regulatory proteins could promote cell damage and bacterial dissemination, while interference with the heat shock protein β1, α-enolase, and the ubiquitin–proteasome system may compromise host control of intracellular *S. aureus*.

Using a screen of recombinant peptides, Dubin and colleagues (19) demonstrated a strong preference of SplB for the WELQ motif (P4–P1) (19). However, none of the natural substrates identified in this or in previous studies (21)(22) contained exactly this cleavage motif. While tryptophan (W) at P4 and glutamic acid (E) or leucine (L) at P3/P2 were occasionally observed, no consistent consensus motif emerged beyond Q at P1. This is in line with the cleavage sites reported in the complement factors C3, C4 and C5 (21). Notably, although Q was highly conserved at the P1 position, cleavage by SplB was highly selective. Not every Q within a protein was cleaved, and, of course, not every protein containing Q was cleaved. As shown in Figure 1A, SplB^WT^ failed to cleave serpin A12 and the sodium/potassium-transporting ATPase subunit α-2, even though they contain numerous Q residues in their sequences. Hence, the requirements for protein cleavage by SplB extend beyond a simple Q at P1 or a short amino acid consensus sequence, instead, an appropriate structural context within the protein is needed.

In conclusion, we identify SplB as a multifunctional protease with a unique substrate spectrum: The enzyme cleaves the heat shock proteins β1, intermediate filaments, α-enolase, and immunoglobulins. Notably, some of these substrates are strictly intracellular (e.g., desmin, vimentin, nestin), while others such as IgG are extracellular, underscoring the capacity of SplB to act across distinct host compartments. This diverse substrate range suggests that SplB contributes to pathogenesis both during the extracellular and intracellular phases of the *S. aureus* lifestyle by destabilizing cytoskeletal scaffolds, impairing stress recovery as well as interfering with metabolic and immune functions. Together, these novel substrates and activities highlight SplB as a versatile factor in the interplay of *S. aureus* with its human host.

## Notes

### Competing Interest Statement

The authors have declared no competing interest.

