## Supplementary Figure and Tables for "Identification of novel human cellular substrates of *Staphylococcus aureus* serine protease SplB"

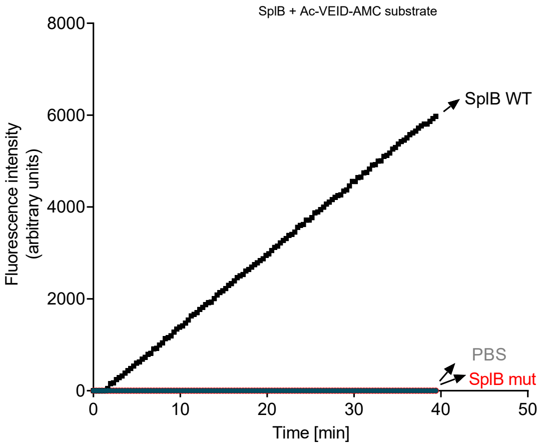


**Supplementary figure 1**: 2.5 µM SplB WT cleaving 25 µM of Ac-VEID-AMC (adapted from Dubin et al., 2008). n = 3 (technical replicates). Abbreviations: amino methyl-coumarin (AMC), acetyl (AC), valine (V), glutamate (E), isoleucine (I), aspartate (D). Measurements were performed in triplicates.

supplemental table 1

### Reversed phase liquid chromatography (RPLC)

| ***Instrument*** | ***Ultimate 3000 RSLC (Thermo Scientific)*** | |
|---|---|---|
| *Trap column* | 75 µm inner diameter, packed with 3 µm C18 particles (Acclaim PepMap100, Thermo Scientific) | |
| *Analytical column* | Accucore 150-C18, (Thermo Fisher Scientific)<br>25 cm x 75 µm, 2,6 µm C18 particles, 150 Å pore size | |
| *Buffer system* | binary buffer system consisting of 0.1% acetic acid in HPLC-grade water (buffer A) and 100% ACN in 0.1% acetic acid (buffer B) | |
| *Flow rate* | 300 nl/min | |
| *Gradient* | *HUNTER samples*<br>0 min 2% B →<br>2 min 5% B →<br>10 min 7% B →<br>40 min 25% B →<br>45 min 40% B →<br>47 min 90% B →<br>53 min 90% B →<br>55 min 2% B →<br>65 min 2% B | *Cross-linking samples*<br>0 min-2% B →<br>2 min-5% B →<br>10 min-7% B →<br>70 min-25% B →<br>75 min-40% B →<br>77 min-90% B →<br>83 min-90% B →<br>85 min-2% B →<br>95 min-2% B |
| *Gradient duration* | 65 min | 95 min |
| *Column oven temperature* | 40°C | |

supplemental table 2

HUNTER - Mass spectrometric analyses_data dependent analyses (DDA)

| ***instrument*** | ***ExplorisTM 480*** |
|---|---|
| *operation mode* | data-dependent |
| ***Full MS*** | |
| *MS scan resolution* | 60,000 |
| *RF Lens (%)* | 40 |
| *Normalized AGC target* | 300% |
| *Microscans* | 1 |
| *maximum ion injection time for the MS scan* | Auto |
| *Scan range* | 350 to 1200 m/z |
| *polarity* | positive |
| *Spectra data type* | profile |
| ***Filter*** | |
| *MIPS* | MIPS Mode=proteins, relaxed restriction when too few Precursors are found=True |
| *dynamic exclusion* | exclude after n times =1<br>Exclusion duration (s) =20, exclude isotopes = True<br>Mass tolerance low=10, high=10 |
| *Minimum Intensity* | 5000 |
| *Charge State* | 2-6 |
| ***Data dependent properties*** | |
| *Number of dependent scans* | 20 |
| ***dd-MS2*** | |
| *Resolution* | 15,000 |
| *MS/MS AGC target* | Standard |
| *maximum ion injection time for the MS/MS scans* | 22 ms |
| *First mass (m/z)* | 22 |
| *Spectra data type* | centroid |
| *isolation window* | 1,4 *m/z* |

| | |
|---|---|
| *Fixed first mass* | 110 m/z |
| *normalized HCD collision energy* | 30% |
| *RF Lens (%)* | 50 |

supplemental table S3

Cross linking - Mass spectrometric analyses_data dependent analyses (DIA)

| ***instrument*** | ***Orbitrap Exploris 480*** |
|---|---|
| *electrospray* | Nanospray Flex™ Ion Source |
| *operation mode* | data-independent |
| ***Full Scan Properties*** | |
| *MS scan resolution* | 120000 |
| *AGC target* | 3e6 (300%) |
| *maximum ion injection time for the MS scan* | 60 ms |
| *Scan range* | 350 to 1200 m/z |
| *Microscans* | 1 |
| *Polarity* | positive |
| *RF Lens* | 50% |
| *Spectra data type* | profile |
| ***Dia Properties (MS2)*** | |
| *Resolution* | 30,000 |
| *maximum ion injection time for the MS/MS scans* | auto |
| *Normalized AGC target* | 3E6 |
| *Spectra data type* | profile |
| *Microscans* | 1 |
| *isolation window* | 66 |
| *Isolation window width* | 13 m/z |
| *Window overlay* | 2 m/z |
| *Fixed first mass* | 200 |
| *HCD collision energy* | 30% |

supplemental table S4

Enrichment ratios of substrate candidates in the cross-linking experiment performed once with SplB wild-type and once with the SplB mutant. *Ctl*: control group without SplB; *1x*: group with 10 µg/mL SplB; *2x*: group with 20 µg/mL SplB. Substrate candidates showing an enrichment factor (2x/1x) >1.2 and absent in the control group (NA) were selected for cleavage testing.

| | Cross linking with SplB mutant | | | Cross linking with SplB Wild-Type | | |
|---|---|---|---|---|---|---|
| Substrate name | 1x/ctl | 2x/cl | Enrichment ratio (2x/1x) | 1x/ctl | 2x/cl | Enrichment ratio (2x/1x) |
| DESM_HUMAN | NA | NA | 2.73 | NA | NA | NA |
| HSPB1_HUMAN | NA | NA | 2.070 | NA | 2.578 | NA |
| IGKC_HUMAN | NA | NA | 1.215 | NA | NA | NA |
| SPA12_HUMAN | NA | NA | 2,448 | NA | NA | 2.77 |
| ENOA_HUMAN | NA | NA | 2,410 | NA | NA | 0.93 |
